# Plasma cells in human pancreatic ductal adenocarcinoma secrete antibodies to self-antigens

**DOI:** 10.1101/2023.03.20.533453

**Authors:** Min Yao, Jonathan Preall, Johannes Yeh, Darryl Pappin, Paolo Cifani, Yixin Zhao, Sophia Shen, Philip Moresco, Brian He, Hardik Patel, Amber Habowski, Daniel A. King, Kara Raphael, Arvind Rishi, Divyesh Sejpal, Matthew Weiss, David Tuveson, Douglas Fearon

## Abstract

Intratumoral B cell responses are associated with more favorable clinical outcomes in human pancreatic ductal adenocarcinoma (PDAC). However, the antigens driving these B cell responses are largely unknown. We sought to discover these antigens by using single-cell RNA sequencing (scRNA-Seq) and immunoglobulin (Ig) sequencing of tumor-infiltrating immune cells from seven primary PDAC samples. We identified activated T and B cell responses and evidence of germinal center reactions. Ig sequencing identified plasma cell (PC) clones expressing isotype-switched and hyper-mutated Igs, suggesting the occurrence of T cell-dependent B cell responses. We assessed the reactivity of 41 recombinant antibodies that represented the products of 235 PCs and 12 B cells toward multiple cell lines and PDAC tissues, and observed frequent staining of intracellular self-antigens. Three of these antigens were identified: the filamentous actin (F-actin), the nucleic protein, RUVBL2, and the mitochondrial protein, HSPD1. Antibody titers to F-actin and HSPD1 were elevated in the plasma of PDAC patients, and also detectable in healthy donors. Thus, PCs in PDAC produce auto-antibodies reacting with intracellular self-antigens, which may result from promotion of pre-existing, autoreactive B cell responses. These observations indicate that the chronic inflammatory microenvironment of PDAC can support the adaptive immune response.

## INTRODUCTION

Microsatellite-stable PDACs do not respond to immunotherapy, have few T cells in cancer cell nests, and have been thought to minimally stimulate the adaptive immune system (*1*). Recently, however, it has been observed that PDAC patients whose tumors contained tertiary lymphoid structures (TLS) with germinal centers, memory B cells, and memory CD4+ T cells had improved long-term survival (*2-4*), suggesting the occurrence of clinically relevant, ongoing anti-PDAC immune responses. The possibility of these immune responses is supported by a study involving patients with PDAC and colorectal cancer in which one-week continuous administration of an inhibitor to the chemokine receptor, CXCR4, revealed on-going anti-tumor immune responses (*5*).

The antigens driving these intratumoral immune reactions in human PDAC are unknown. Although PDAC has a low mutational frequency compared to other cancers (*1*), attention has been directed to T cell clones from PDAC patients with specificity to neoantigens arising from mutations that predicted cross-reactive microbial epitopes (*6*). The intratumoral immune response may also be directed towards germline-encoded antigens, a concept that has been supported by the finding that immunization of mice with induced pluripotent stem cells confers protection against several tumor models (*7, 8*). Thus, defining the range of antigens that are driving the intratumoral immune response in PDAC may expand our knowledge of the interaction between this cancer and the immune system.

B cell activation is triggered by antigen interaction with membrane immunoglobulin (Ig). This interaction may lead to B cell activation, antigen-presentation to CD4 T cells, and the formation of germinal center reactions. The latter will lead to B cell clonal expansion, heavy (H) chain isotype switching, H chain and light (L) chain variable region somatic hypermutation (SHM), and ultimately differentiation into antibody-secreting PCs. Therefore, antibodies derived from activated B cell responses may help identify those antigens that are driving intratumoral immune responses. In the current report, we have used scRNA-Seq of PCs and B cells from primary PDAC specimens to identify their paired H and L chains which enabled screening of antibody reactivity.

## RESULTS

### Identification of B and T cell responses by scRNA-Seq in human PDAC

We performed scRNA-Seq of CD45+ immune cells from seven untreated primary PDAC samples, including four microsatellite-stable surgically resected tumors and three fine-needle aspiration (FNA) biopsies (Table S1). scRNA-Seq identified a total of 26,702 immune cells and 327 non-immune cells. The major immune cells included T cells (66%), myeloid cells (19%), PCs (7.4%) and B cells (3.6%), with T and myeloid cells being the most frequent cell types in each sample (Fig. 1A-B and Fig. S1A). In the *CD8+* T cells, features of T cell activation were prominent, as evidenced by frequent expression of the effector genes, *PRF1* (46% of CD8 T cells) and *GZMB* (31%), as well as the cytokines, *IFNG* (13%) and *TNF* (11%). Subsets of *CD8*+ T cells also expressed the inhibitory receptors, *PD-1* (14%) and *CTLA4* (3%) (Fig. 1C and Fig. S1B), which also identify effector cells. In the *CD4+* T cell population, a Th1 response was apparent, as assessed by the expression of the Th1 lineage marker, *TBX21* (13% of CD4 T cells), and the Th1 cytokines, *TNF* (18%) and *IFNG* (6%). *FOXP3+ CD4+* regulatory T cells represented a relatively abundant CD4 T cell population (22%), and characteristically co-expressed *IL2RA*, *PD-1* and *CTLA4*. *CD4+* T follicular helper T cells (Tfh cells) were identified by their expression of *BCL6* (2%), *IL-21* (1%), and the chemokine receptor, *CXCR5* (2%) (Fig. 1C and Fig. S1C-D). In summary, the PDAC tumors contained activated CD8 and CD4 T cells, including Tfh cells of presumed germinal center origin.

**Fig. 1.**
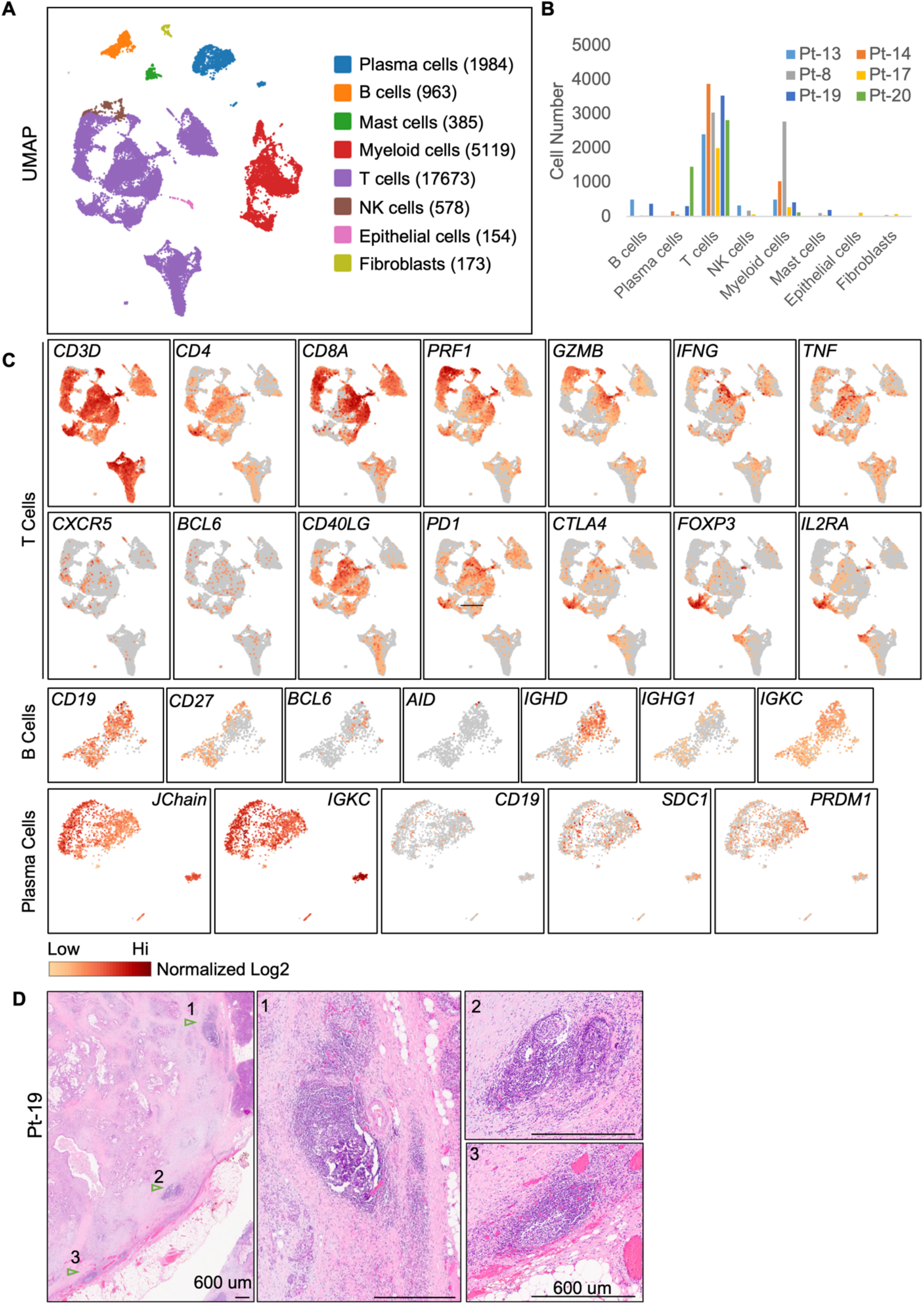
Identification of active T and B cell responses in PDAC by scRNA-Seq. (**A**) UMAP clusters are shown of immune cells isolated from six primary PDAC. (**B**) The cell numbers of different immune cell types identified in individual patient are shown. (**C**) The expression of selected genes in T cell, B cell and PC clusters are shown. The intensity of the colors represents the normalized log2 transformation of UMI. (**D**) Examples of TLS like structures (indicated by arrowhead and labelled with 1,2 and 3) in patient-19. The zoom in examples of TLS were shown in the right. Scale bar represents 600um.

The B cell cluster (n=963 cells) was defined by the expression of the pan B cell markers, *CD19* and *CD20*. 35% of B cells expressed the naïve H chain isotype, *IgD*, and B cells that expressed isotype switched *IgG1* (19%) were also present (Fig. 1C). Additional evidence for an active intratumoral B cell response was indicated by the presence of germinal center B cells, as assessed by their expression of *BCL6* (4%) and *AICDA* (1%), the latter of which mediates SHM and isotype switching (Fig. 1C and Fig. S1E). PCs (n=1984) were identified by the makers, *J chain*, *PRDM1*, and *CD138*, and high expression of Ig H and L chains (Fig. 1C). Most PCs (95% of PCs) no longer expressed *CD19*, indicating they were relatively mature PCs rather than plasmablasts. Thus, PDAC tumors contained activated B cells and terminally differentiated PCs.

We further obtained histopathology images from the four resected PADC samples which we have performed scRNA-Seq. We have observed presence of multiple TLS-like structures in each tumor sample, with TLSs typically located in the tumor border (Fig. 1D and Fig. S2). This observation supports that the B and T cell response revealed by scRNA-Seq may take place inside those TLSs.

### Igs from PCs in PDAC featuring isotype switching, somatic hypermutation and clonal expansion

A total of 615 PCs with paired H and L chains were detected in six PDAC samples (Fig. 2A and Fig. S3A). A minority of PCs (0-17%) expressed the *IgM* isotype, while the majority had undergone class switching to *IgG1*, *IgG2*, *IgA2*, or *IgA1* isotypes (Fig. 2B and Fig. S3B). Analyzing SHM in the V regions revealed an average of 23 and 15 mutations in the H and L chains, respectively (Fig. 2C). These rates of SHM are comparable to those in antibodies induced by viral infections (*9*). Expanded PC clones were identified in four of the six PDAC samples, with clone sizes ranging from 2 to 46. Strikingly, in patients 19 and 20, more than half of the Ig-paired PCs were products of clonal expansion (Fig. 2D). Analysis of the evolution of V region SHM sequences revealed that most PCs originated from single expanded nodules, although additional lineage evolutions were present (Fig. 2E and Fig. S3C), consistent with typical germinal center reactions. Fewer B cells with paired Igs were sequenced (n=473), likely due to lower Ig expression, which yielded only four small, expanded clones (clone size 2-3). Therefore, PCs in PDAC likely arose from germinal center reactions in the PDAC stroma.

**Fig. 2.**
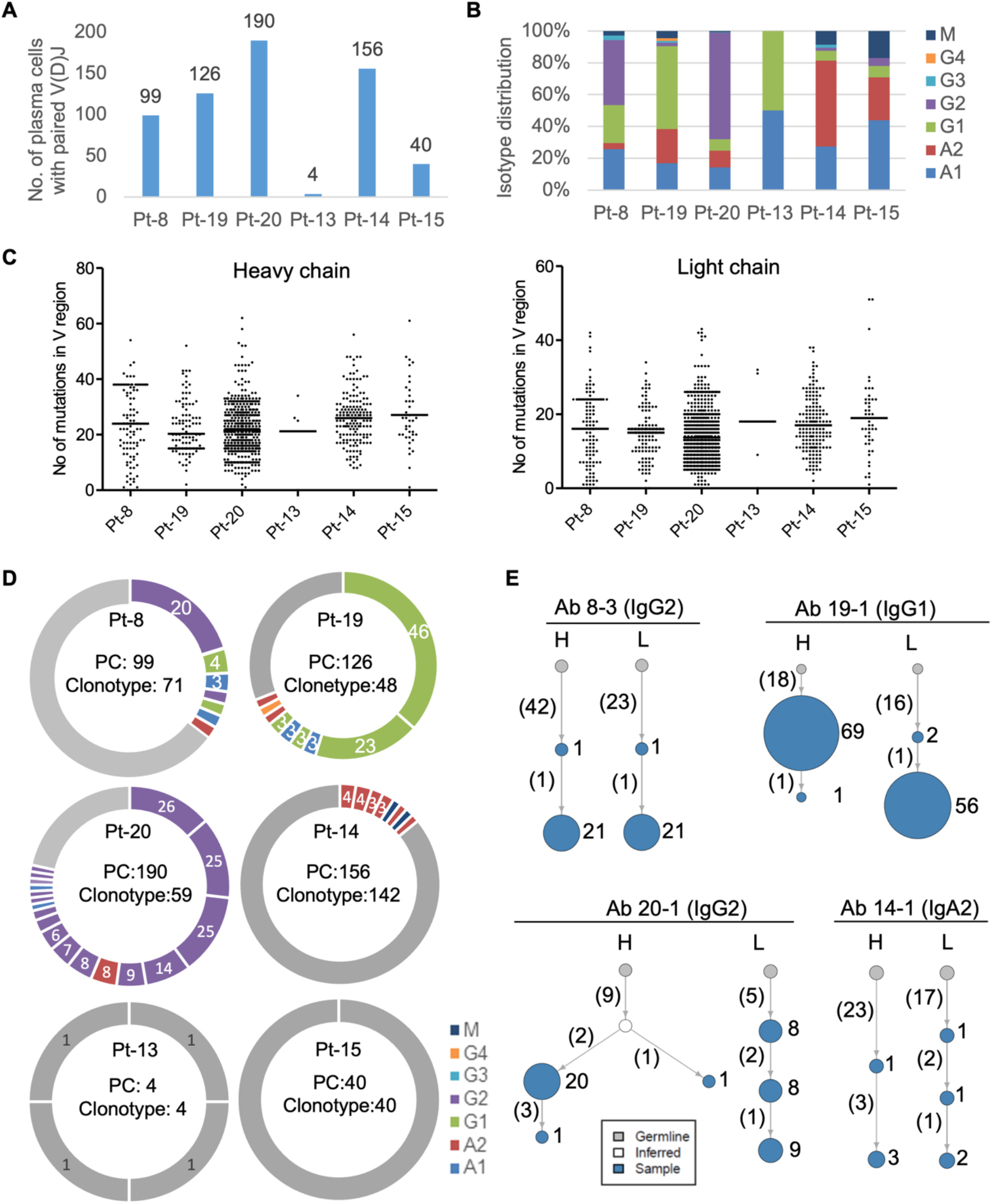
Identification of isotype switched and somatically hypermutated Igs by sequencing of PCs from PDAC. (**A**) Numbers are shown of PCs in which paired H and L chains were sequenced in each patient. The distribution of Ig isotypes (**B**) and the number of somatic mutations in V regions (**C**) in PCs from individual patients are determined. (**D**) The PCs with paired H and L chains and their clonotype distribution are shown for each patient. PCs within expanded clones are labelled with colored doughnuts, with colors indicating isotypes. Each slice represents one clone, with clone size proportional to slice size. Single clone PCs are pooled and labeled with gray color. (**E**) Examples are presented of antibody lineage evolution within expanded PC clones for individual patients. Clone size is indicated by the size of the node (not scaled) and labeled by numbers on the right. The number of somatic mutations in the combined V(D)J regions is shown in parenthesis.

### Antibodies from PCs binding to intracellular self-antigens

Forty-one antibodies with paired H and L chains from the six PDAC patients were selected for recombinant antibody synthesis. These antibodies included the mostly expanded PC cell clones, as well as some single-clone PC and B cells, and represented the products of 235 PCs and 12 B cells (Fig. 3A and Table S2). Immunofluorescent staining of human PDAC cell lines and tumors was used to screen the reactivity of the recombinant antibodies. Twenty-five of the 41 antibodies showed positive binding to PDAC cell lines, with the expanded clones showing more frequent binding (Fig. 3A). Recombinant antibodies reacted with antigens in all sub-cellular locations, including cytoplasmic (e.g. 8-3 and 15-7), cytoplasmic-enriched (e.g. 19-1), both cytoplasmic and nuclear (e.g. 19-4), and nuclear-enriched reactivities (e.g. 19-3 and 20-1) (Fig. 3B-C and Fig. S4). We did not identify any antibodies staining cell surface antigens. Antibodies derived from a single patient’s PDAC tumor could exhibit diverse staining patterns (for example, 19-1 and 19-3), indicating the occurrence of an adaptive immune response to multiple intracellular antigens in the same patient. All the reactive antibodies could stain multiple PDAC cell lines with a similar staining pattern, as well as non-tumor human pancreatic ductal cell line, HPDE, and human fibroblasts (Fig. 3D and Fig. S5). When staining PDAC tumors from which antibody sequences were derived, the antibodies bound both to cancer cells and stromal cells (Fig. 3E). Thus, non-mutated, non-cancer cell-specific intracellular antigens drive common humoral immune responses in human PDAC.

**Fig. 3.**
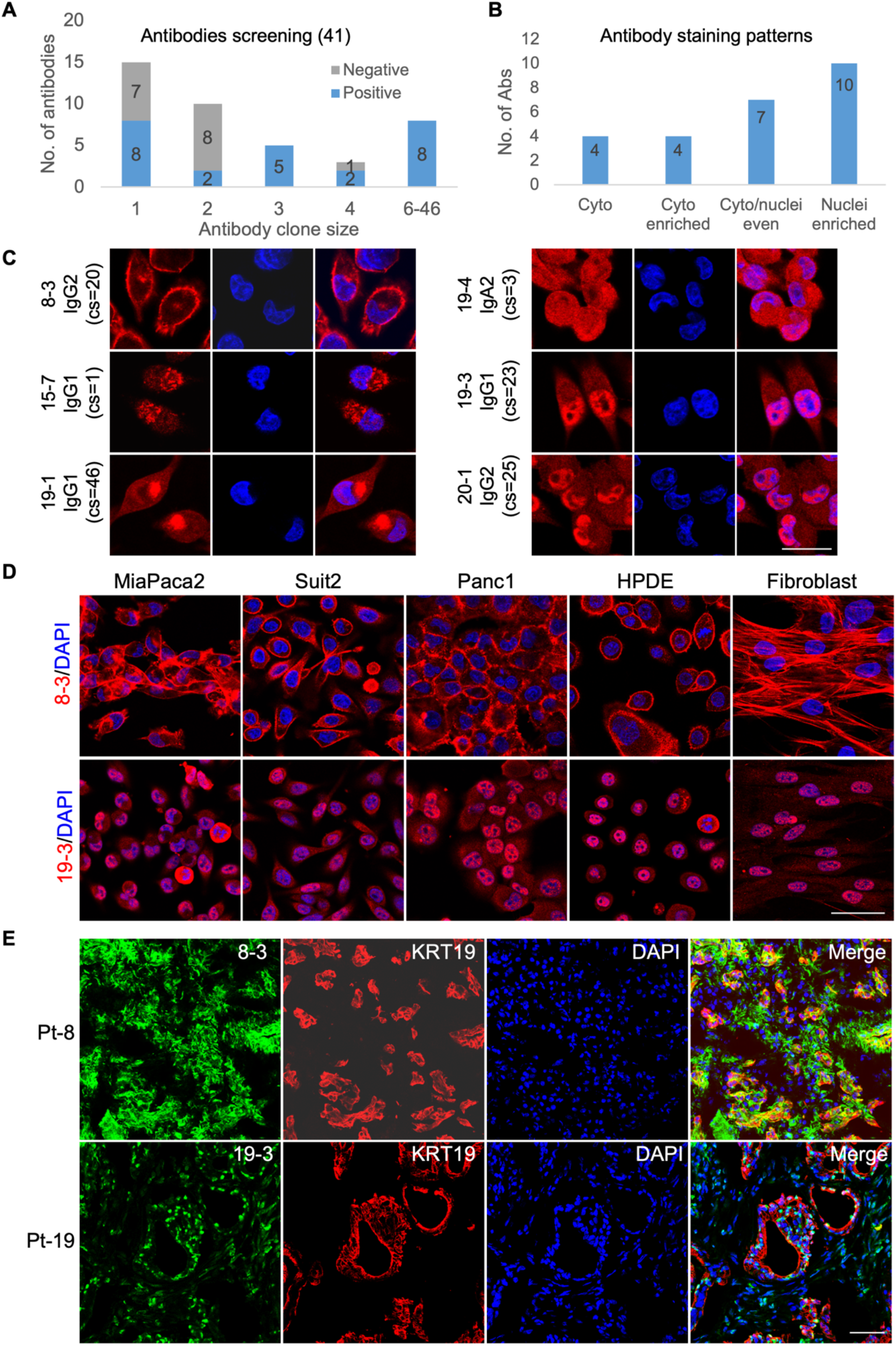
Recognition of intracellular self-antigens by recombinant antibodies from expanded PCs of PDAC tumors. (**A**) Summary is presented of the results of staining various cell lines by 41 recombinant antibodies based on scRNA-Seq of PCs and B cells from six PDAC tumors, distributed according to clone sizes. (**B-C**) The distribution and examples are shown of four distinct antibody staining patterns: cytoplasmic (Cyto, e.g. 8-3 and 15-7), cytoplasmic enriched (Cyto enriched. e.g. 19-1), evenly distributed in cytoplasmic and nuclei (Cyto/nuclei even, e.g. 19-1), and nuclei enriched (e.g. 19-3 and 20-1). Examples of positive antibody staining (red) in MiaPaca2 cell line, co-stained with DAPI (blue). Original antibody isotype and clone size (cs) are indicated. (**D**) The recombinant antibodies, 8-3 and 19-3 (red), respectively, were used for staining different PDAC and non-cancer cell lines, along with nuclear staining with DAPI (blue). (**E**) Antibodies 8-3 and 19-3, respectively, were used to stain the PDAC tumors from which their PCs were derived. Staining was also done with anti-KRT19 antibody to demonstrate the cancer cells. Scale bar in C, 25um, and in D and E, 50um.

### Identification of antigens driving PC cell response in PDAC

Three PDAC antigens were identified by using antibody-mediated immunoprecipitation from MiaPaca2 cell lysates, followed by mass spectrometry analysis. Antibody 8-3, which was derived from the most expanded PC clone (clone size 20) from patient 8, recognized a cortical structure in MiaPaca2 cells (Fig. 3C). This antibody immunoprecipitated a major protein band of approximately 45 kDa, which was identified as ACTIN by mass spectrometry, as well as the ACTIN associated proteins, MYH10 and MYH9 (Fig. 4A). Antibody 8-3 co-localized with filamentous actin (F-actin) in the cell cortex, and its antigen re-localized to perinuclei foci after treatment with the actin destabilizing drug, cytochalasin D (Fig. 4B). Knockdown of ACTIN with siRNA reduced staining by antibody 8-3, but not staining by a MYH10-specific antibody (Fig. S6A), indicating that ACTIN was the target of antibody 8-3. This antibody was confirmed to bind to polymerized F-actin, but not monomeric G-actin, using in vitro actin polymerization and depolymerization assays (Fig. 4C). We developed an F-actin specific ELISA and measured the EC50 of antibody 8-3 for F-actin to be 9.1 nM, a relatively high affinity/avidity that was probably a result of SHM (Fig. 4D and Fig. 2E). Sequencing the actin genes (ACTB and ACTG1) from the tumor organoid derived from patient 8, the source of the PCs encoding antibody 8-3, showed no mutations in the protein-coding regions, confirning the self-antigen nature.

**Fig. 4.**
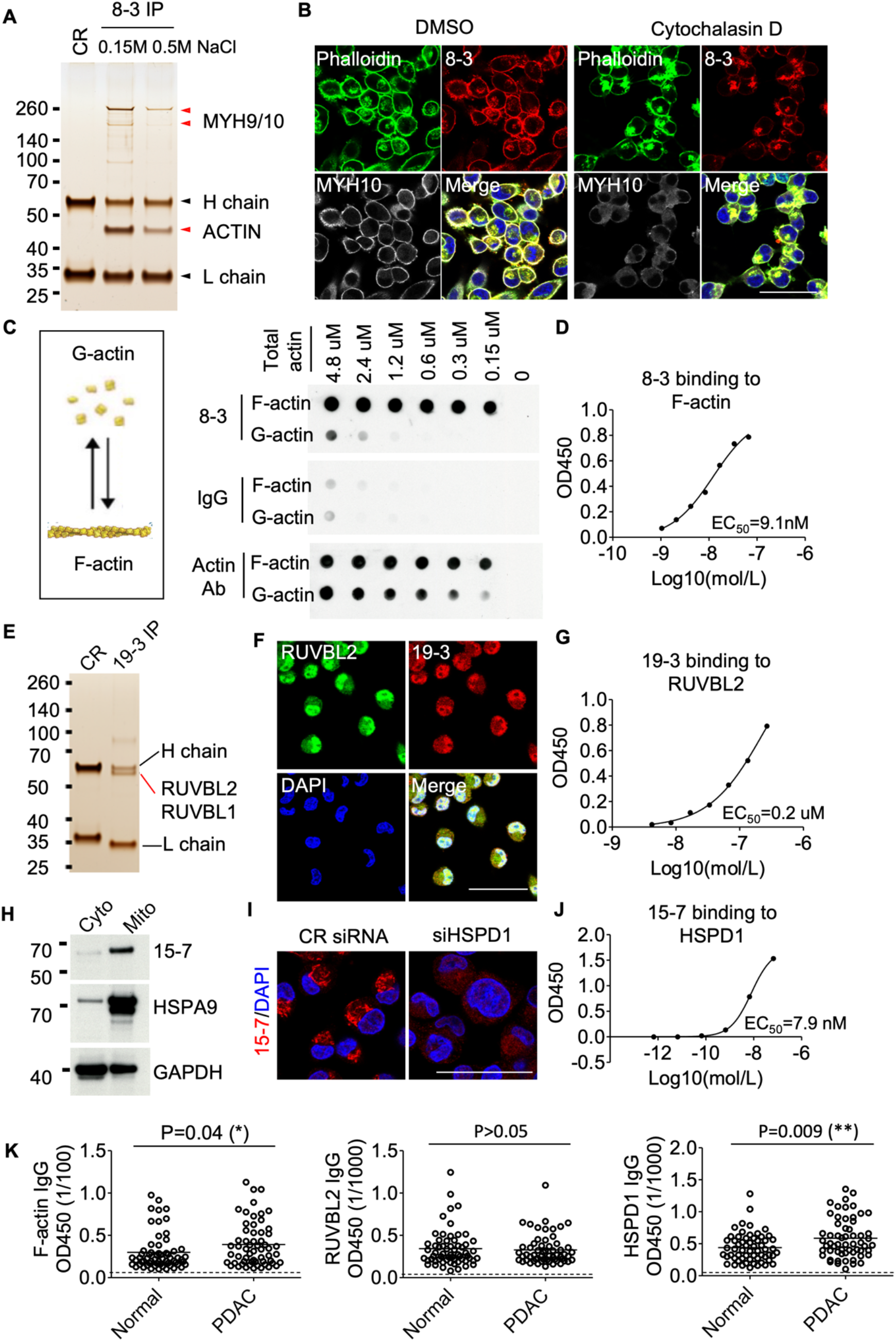
Identification of F-actin, RUVBL2 and HSPD1 as antigens in PDAC. (**A**) Proteins that were immunoprecipitated by antibody 8-3 from lysates of MiaPaca2 cells were separated by SDS-PAGE and visualized by silver staining. Two bands were identified by mass spectrometry (red arrowheads). (**B**) MiaPaca2 cells were co-stained with 8-3, phalloidin, and an antibody specific for MYH10 after 30min treatment of the cells with DMSO or cytochalasin D. (**C**) Dot-blot assay is shown the reaction with F-actin or G-actin of 8-3, isotype control IgG, and a commercial actin specific antibody. (**D**) The binding of incremental concentrations of 8-3 to F-actin was measured by ELISA. (**E**) Proteins that were immunoprecipitated by recombinant antibody 19-3 from lysates of MiaPaca2 cells were separated by SDS-PAGE and visualized by silver staining. The proteins, RUVBL1 and RUVBL2, were identified by mass spectrometry. (**F**) MiaPaca2 cell were co-stained with 19-3 and an antibody specific for RUVBL2. (**G**) The binding of incremental concentrations of 19-3 to RUVBL2 was measured by ELISA. (**H**) Western blot analysis of recombinant antibody 15-7 detection of cytoplasmic proteins (Cyto) and mitochondria-enriched proteins (Mito) is shown. (**I**) Immunofluorescent staining of MiaPaca2 cells by 15-7 and an antibody specific for HSPD1 is shown. (**J**) The binding of incremental concentrations of 15-7 to HSPD1 was measured by ELISA. (**K**) The serum IgG titers of normal individuals and PDAC patients to F-actin (sera diluted 1:100), to RUVBL2 (sera diluted 1:1000), and to HSPD1(sera diluted 1:1000), respectively, were measured by ELISA. The non-parametric t-test was used for comparison. Scale bars in **B**, **F** and **I** are 50um.

Similarly, antibody 19-3, the second most expanded PC clone in patient 19 (clone size 23), immunoprecipitated a protein complex of RUVBL1 and RUVBL2, as determined by mass spectrometry. Knockdown of RUVBL2 reduced antibody 19-3 binding in the MiaPaca2 cell line (Fig. 4F and Fig. S6B). This antibody recognized recombinant RUVBL2 but not RUVBL1 protein, with an EC50 of 0.2 uM (Fig. 4G and Fig. S6C), confirming that antibody 19-3 recognizes RUVBL2.

Antibody 15-7, which was derived from a single PC clone in the tumor from patient 15, co-stained with antibodies specific for the mitochondrial markers, COX4I1 and HSPA9 (Fig. S6D). This antibody recognized a mitochondrial-enriched protein band with an apparent molecular weight of 60 kDa (Fig. 4H), which was identified as heat-shock protein D1 (HSPD1) by mass spectrometry. HSPD1 was confirmed to be the antigen of antibody 15-7 by immunofluorescent co-localization in MiaPaca2 cells, siRNA knockdown, and recombinant protein binding, with an EC50 of 7.9 nM (Fig. 4I-J and Fig. S6D).

In summary, our identification of three self-antigens, including antigens for two antibodies from highly expanded PC clones, indicates that widely expressed, intracellular self-antigens may be targets of humoral immune responses in human PDAC.

### The frequent occurrence of antibody responses to F-actin and HSPD1 in PDAC patients

We collected plasma samples from 59 PDAC patients and 61 healthy donors, and measured IgG titers to F-actin, RUVBL2, and HSPD1, respectively (Table S3). Demographics, such as gender, age, and race, did not significantly affect antibody concentration. Interestingly, plasma from normal donors contained detectable levels of IgG responses to these three antigens. In PDAC, the IgG F-actin and HSPD1 antibody titers were significantly higher than those in normal individuals (Fig. 4K and Fig. S7A). IgG F-actin antibody titers were significantly reduced in patients who had received neoadjuvant therapy, but this was not observed for either HSPD1 or RUVBL2 antibody titers (Fig. S7B), suggesting that B cell responses for different antigens may have different sensitivities to cytotoxic chemotherapy. These antibody responses did not correlate with other pathological features such as tumor stage, size or grade. Thus, PDAC patients likely promote auto-antibody responses which pre-exist in healthy donors.

## DISCUSSION

PDAC is generally considered to be one of the more immunologically “silent” carcinomas and to be resistant to cancer immunotherapy for this reason. Reports, however, of TLSs in PDAC (*2-4*), of neoantigens with homology to infectious disease-derived peptides in long-term survivors of PDAC (*6*), and of signs of improved anti-PDAC immunity in metastatic lesions of patients with PDAC after inhibition of CXCR4 (*5*), all point to a need for a deeper understanding of the relationship between this cancer and the immune system. The antigens driving the B cell response are more readily identified by the Ig product of the B cells and PCs. Thus, we chose to interrogate the Ig products of intratumoral B/PCs in PDAC.

Our scRNA-Seq analysis of CD45+ immune cells in primary PDAC samples indicated active intratumoral immune reactions, such as the presence of effector CD8 T cells expressing *PRF1*, *GZMB*, and *IFNG*, and effector CD4 T cells expressing *TNF* and *IFNG*. We were able to confirm the four resected PDAC tumor samples as being microsatellite-stable, representing the majority of PDAC (*10*). The presence of effector T cells in our cohort is consistent with previous studies of human and mouse autochthonous PDAC immune microenvironments in which activated, effector T cells were found (*11*). In addition, scRNA-Seq identified CD4 T cells with the Tfh phenotype by their transcriptional expression of *BCL6* and *CXCR5*, and germinal center B cells expressing *BCL6*. The presence of a B cell response is further supported by Ig sequencing that identified isotypically switched, somatically hypermutated and expanded PC clones. The presence of TLS in PDAC stroma from the paired histology images further support the integrated B and T cell responses. It is interesting that these indicators of an active adaptive immune reaction are occurring despite the presence of large proportion of FOXP3-expressing regulatory T cells.

There are a few previous studies exploring the antigen reactivity of B cells and PCs in other tumors (*12, 13*), but our study is the first to apply this approach to PDAC. While we were unsuccessful in our initial intent to discover tumor-specific B cell antigens, the finding of Igs from PCs frequently binding to widely expressed, non-mutated intracellular self-antigens in human PDAC is intriguing. Our screening in PDAC identified most antibodies as recognizing intracellular antigens that were shared by multiple PDAC cell lines, normal epithelial cells and fibroblasts. This conclusion was confirmed by the identification of three self-antigens, F-actin, RUVBL2 and HSPD1, as targets of these humoral immune responses. All the seven PDAC patients enrolled for this study have no records of auto-immune disease or pancreatitis. In this context, it is worth noting that 20% of naïve B cells in human peripheral blood are reported to be autoreactive (*14*), which is consistent with our detection of autoantibody responses to those antigens in healthy donors. Self-reactive B cell responses are likely promoted in PDAC because of continual exposure of intracellular self-antigens through cell death and chronic inflammation, reminiscent of the antibody responses in autoimmune diseases. Autoantibody responses to F-actin, RUVBL2 and HSPD1 have been reported in various autoimmune conditions (*15-17*) and cancers (*18, 19*). Previously, immunization with mouse syngeneic ES cells reduce mouse pancreatic tumor progression, indicating self-antigens likely drive such immune response (*7, 8*). Additionally, a recent study reported that the self-protein matrix metallopeptidase 14 was a major auto-antigen in human ovarian cancer, and some Igs were likely originated from germline encoded auto-reactive B cell (*13*). Thus, the B cell response to self-antigens may be a common feature of cancer.

Our analysis of plasma titer to F-actin, RUVBL2 and HSPD1 did reveal that PDAC patients have significantly higher titers to F-actin and HSPD1 as compared to a group of healthy donors. Our analysis did not reveal a significant correlation with tumor-associated pathological factors, although this may be partially due to our relatively small cohort of patients. Our studies can be potentially strengthened by analyzing serial tumor biopsies and plasma obtained during the course of tumor progression or response to therapy such as CXCR4 inhibition, thus providing more dynamic pictures of B cell response and antibody titer changes.

One open question from this study is whether the B cell response to intracellular self-antigens affects PDAC progression. Given the generally positive correlation of a B cell response with a favorable outcome in PDAC (*2-4*), we speculate that those autoreactive B cell responses may indirectly have an anti-tumor role. While antibodies binding intracellular antigens cannot directly target live tumor cells, these antigens can be exposed to antibody binding during cancer cell death, such as necrosis, and potentially induce inflammation through complement activation and Fc receptor binding. Moreover, we expect that B cells binding such self-antigens will internalize and engage in antigen presentation to CD4+ T cells. F-actin has been reported to promote antigen presentation through the F-actin receptor Clec9A on dendritic cells in breast cancer model (*20*). Further understanding the B cell response in PDAC may potentially provide opportunities for modulation of cancer immunotherapy. In any event, the finding of T cell and B cell responses in the PDAC tumors indicates that an adaptive immune response is occurring in the tumor microenvironment, which may be exploited for tumor control.

## MATERIALS AND METHODS

### Study design

This study is designed to study B cell response and identify B cell targeted antigens in human PDACs. This study consists of: 1) ScRNA-Req of immune cells infiltrating human PDAC and Igs of B and plasma cells; 2) Synthesizing of recombinant antibodies from Ig sequencing; 3) Screening of recombinant antibodies reactivities toward PDAC cell lines and tumor tissues; 4) Identification of antigens. This pipeline was used to study four resected and three FNA naïve-treated PDAC samples. The antibodies synthesizing and screening aim to profile the most expanded plasma cells clones in PDAC to get a better picture of B cell reactivity in human PDAC.

### Human samples

All human samples were obtained with written consent and IRB approval. Four resected PDAC specimens and three FNA (two passes each) specimens were obtained from Northwell Health. Archived plasma samples from PDAC patients (n=59) were received from Northwell Health. Plasma samples from healthy donors (n=61) were obtained from volunteer donors or purchased from commercial resources (Innovative Research and BioChemed Services). Microsatellite instability status of resected PDAC samples were determined by immunohistochemistry staining of MLH1, MSH2, MSH6 and PMS2, performed and interpreted by pathologists from Northwell Health. Histopathologic hematoxylin and eosin images were obtained from Northwell pathology department acquired by Leica Aperio slide scanner. All patient samples were de-identified.

### Cell lines and culture

Human PDAC cell lines MiaPaca2, Suit2, Panc-1, and foreskin fibroblast (BJ-hTert) were obtained from Cold Spring Harbor Laboratory cell validation center, and had been validated by short tandem repeat profiling. The normal human pancreatic ductal epithelial cell line (HPDE) was obtained from Dr. Krainer lab in CSHL, originally purchased from Kerafast. All cell lines had been tested mycoplasma free and cultured in the conditions recommended by ATCC or the manufacturer.

### Tumor samples preparation and Single-cell sequencing

A small portion of the resection tumor was snap-frozen for histology. The remaining human PDAC tumor samples were digested into single cells using collagenases and about 10% of single cells digest was used for tumor organoid culture as described (*21*). Live immune cells were sorted by EpCAM^-^ /CD45^+^ (Biolegend, # clone 2D1 and 9C4) on a Sony SH800 cell sorter. About 10,000 immune cells were barcoded with 10X Genomics Chromium single cell 5’ kit, and both gene expression and Ig libraries were prepared and sequenced according to the 10X Genomics manual.

### Sequencing data analysis

The scRNA-Seq and Ig reads were aligned and quantified using the Cell Ranger pipeline (10X Genomics, version 6.0.0). Dead cells and cells with the number of genes expressed less than 200 were also filtered out. ScRNA-Seq Data was normalized, logarithm transformed and scaled. Doublets were removed using a R package ‘Scrublet’. Principle component analysis (PCA) was run with 50 components using the top 4000 genes of each sample. The nearest neighbor algorithm was run with 50 PCAs and clustered in UMAP projections using Leiden clustering. All additional analyses were performed using Loupe Browser (10X Genomics, version 6.1.0) and the Python toolkit Scanpy 1.6.0. Immune cells clusters were defined using the set of markers and UMAP clusters: T cells (CD3^+^, TCR^+^, NCAM^-^), NK cells (CD3^-^, NCAM^+^), B cells (CD20^+^, CD79A^+^, IgKC ^low^), PCs (JChain^hi^, IgKC^hi^), myeloid cells (CD14^+^, CSF1R^+^), mast cells (CPA3^+^, GATA2^+^), fibroblasts (FAP^+^, ACTA2^+^, PDGFRA1^+^), epithelial cells (CK19^+^, CK18^+^, CK5^+^, CK14^+^).

### Ig data analysis

PCs and B cells clusters defined by scRNA-Seq were used to select cell type specific Igs analysis. In one patient (Pt-15), gene expression sequencing data was not available, the PCs were defined by high expression of H chain (UMI>=100). B or PC specific Igs were exported from the Loupe V(D)J Browser (10X Genomics, version 4.0.0) and analyzed for isotypes and clonotypes. For somatic hypermutation analysis, V region sequences were analyzed by Igblast (NCBI) and visualized by Graphpad Prism 5 (Dotmatics). For lineage tree analysis, Ig sequences were first analyzed using R packages Change-O and alakazam (*22*), and the Ig lineages were built using a R package PHYLIP. Finally, Lineage trees were visualized using a R package igraph.

### Recombinant Antibodies production

The selected antibody V(D)J sequences were synthesized from IDT, and cloned into pFUSEss H and L vectors (Invivogen) using NEB HIFI assembly kit (NEB, #5520s). The heavy chains were cloned into human IgG1 or customer-designed human IgG4-His vector. A 6x His tag with a Gly-Ser-Gly linker was inserted before the stop codon of human IgG4 constant region, synthesized and replaced the IgG1 in pFUSE-CHIg-hG1 vector. The light chain was cloned into a vector containing IgKC, IgLC2, or costumed-made IgLC1 vector, to match the original light chain. Sequences confirmed antibody vectors were mixed at H:L 1:1.5 ratio, and transiently transfected into 293T cells, cultured in 10% IgG reduced serum. Culture supernatants were collected four days later and antibodies were purified using the Protein G column (Thermo Fisher, #89956). The antibodies were further concentrated and the buffer was exchanged into PBS using Amicon 50K centrifugal filter device (Sigma, #UFC805096). The antibody concentration was quantified by NanoDrop (Thermo Fisher) and human IgG ELISA kit (Mabtech, #3850-1H-6).

### Antibody screening using immunofluorescent staining

Cells were seeded in the 24-well glass-bottom cell culture plate (Chemglass, #CLS-1812-024), and used 2-3 days later at 50-80% confluency. Cells were fixed with 4% PFA, permeabilized with 0.1% Triton-X100, blocked with 3% FBS, and stained with 10 ug/ml human query antibodies diluted in PBST containing 1% FBS overnight at 4 °C. Cells were then washed three times with PBST, and stained with fluorescent conjugated secondary anti-human IgG antibody (Thermo Fisher, #A21090 and #A-11013) and DNA dye DAPI. In human tumor section staining, anti-His secondary antibody (Biolegend, #362607) was used to detect IgG4-His primary antibodies. Other antibodies and dyes used were: Phalloidin (Thermo Fisher, #A12379), MYH10 (Atlas Antibodies, #HPA047541), cytochalasin D (Thermo Fisher, #PHZ1063), ACTIN (Cell Signaling Technology, #3700S), RUVBL2 (Atlas Antibodies, #HPA067966), RUVBL1 (Atlas Antibodies, # HPA019947), HSPD1 (Atlas Antibody, #HPA050025), HSPA9 (Atlas Antibody, #HPA000898), COX4I1 (Atlas antibody, #HPA002485), KRT19 (Abcam, #ab203445), human IgG isotype controls (Biolegend, #403502 and #403702). The samples were imaged in Leica SP8 confocal microscope under 40x magnification. Antibody screening images were acquired using the same confocal setting for isotype control staining, and the gain of query antibodies was reduced if images were saturated in the setting. Images were analyzed using Leica LAS X or Image J software.

### Immunoprecipitation and mass spectrometry

Antibodies were covalently crosslinked to magnetic beads using the Thermo Fisher Dynabeads Antibody Coupling Kit. The total cell lysate was prepared from MiaPaca2 cell line using TNET lysis buffer (50 mM Tris-HCl, pH 7.5, 150 mM NaCl, 5 mM EDTA and 1% Triton X-100). Dynabeads containing 2-5 ug crosslinked antibodies or isotype control were first blocked with 1%BSA, then incubated with up to 1mg cell lysate overnight in 4C in TNET buffer. The beads were separated using a magnetic rack, washed three times with TNET buffer (or with increased NaCl concentration), and eluted with 2X SDS under boiling. The immunoprecipitation elute was running in gradient SDS-PAGE gel and stained with a silver staining kit (Thermo Fisher, #24600).

Distinct gel bands were excised, de-stained, reduced with 3 mM TCEP and alkylated with 10 mM CEMTS, and then digested with trypsin. Eluting peptides were ionized and transferred into an Exploris Orbitrap mass spectrometer (Thermo Fisher). Spectral data were searched against the human database and a database of common contaminants. M-oxidation and N/Q-deamidation were set as variable modifications. Peptide-spectral matches were filtered to maintain FDR <1% using the Percolator.

### siRNA knockdown

Cells were transfected with control or siRNA at 20-60uM, cultured for three days, and proceeded for immunofluorescent staining. siRNA against human ACTB (#SASI_Hs01_00204238, #SASI_Hs01_00204239), MYH10 (#SASI_Hs01_00072460, #SASI_Hs02_00340636), MYH9 (#SASI_Hs01_00197338, #SASI_Hs01_00197339), RUVBL2 (GATGATTGAGTCCCTGACCAA, GAAGATGTGGAGATGAGTGAG), control (GGATGTAAGTGGGAAAGTGGA) were purchased from Sigma. siRNA targeting HSPD1 (L-010600-00-0005) was purchased from Horizon Discovery.

### Actin polymerization or de-polymerization and dot-blot assay

Human non-muscle actin (Cytoskeleton, # APHL99) was polymerized or de-polymerized according to the manufacturer’s protocol. Briefly, actin was first diluted into 0.4 mg/ml in G-buffer (5 mM Tris-HCl, 0.2 mM CaCl2, 0.2 mM ATP), and kept in ice for 1h. For polymerization, 10% polymerization buffer (500 mM KCl, 20 mM MgCl2) with 1 mM final ATP and 5uM phalloidin (Simga, #P2141) was added to the actin and incubated at room temperature for 1h. For de-polymerization, 5 uM latrunculin B (Sigma, #428020) was added into the actin with G-buffer, and incubated in ice for 1h. Actin samples were further diluted into G-buffer or polymerization buffer supplemented with phalloidin or Latrunculin B, and serial dilutions of F-actin or G-actin were blotted on nitrocellulose membrane using Bio-Dot microfiltration apparatus (Bio-Rad) according to the manufacturer’s instructions. The membrane was furthered blocked and incubated with human IgG4 isotype control, Ab 8-3, mouse anti-human Actin antibody (Thermo Fisher, # MA1-140) at 1 ug/ml for 1h, washed and detected with HRP conjugated anti-human IgG (Thermo Fisher, #A18811) or anti-mouse IgG (Biolegend, #405306) secondary antibody, and developed by ECL substrate.

### ELISA

Recombinant proteins RUVBL2 (Creative Biomart, #RUVBL2-30950), RUVBL1 (Novus, #NBP1-50845), HSPD1 (Origene, #TP760396), or BSA control were coated in PBS with 0.25 ug per well in 96-well ELISA plate (Corning) overnight in 4C. The plate was blocked with 1% BSA, washed with PBST, incubated with query antibodies or isotype control at 1 ug/ml diluted in 0.1% BSA in PBST for 1h, detected with HRP conjugated secondary antibody (1ug/ml), and developed by TMB substrate (Biolegend, #421101) for 5-10 min. For F-actin ELISA, polymerized F-actin was diluted in polymerization buffer supplemented with 5uM phalloidin and 1mM ATP, and coated in the ELISA plate for 1h at room temperature, followed by antibody binding and development.

For human plasma studies, the antigens were coated as above, then blocked with 5% milk in PBS. PDAC or healthy donor plasma was diluted 1/100 and 1/1000 in 0.5% milk in PBST, and each sample was run in duplicates. Positive controls are using antibodies 8-3, 19-3 or 15-7. Background signal was obtained using the higher signal from secondary antibody only and plate coated with BSA. All samples to the same antigen were analyzed at the same time to avoid batch differences. Each ELISA analysis was repeated twice.

### Mitochondria cell fraction and western blot

For crude mitochondria fraction, the cells were first lysed using low salt buffer (0.25 M sucrose, 20 mM HEPES-KOH pH 7.5, 10 mM KCl, 1.5 mM MgCl2, 1 mM EDTA, 1 mM EGTA, 1 mM DTT) supplemented with proteinase/phosphatase inhibitors, homologized with a Teflon-glass homogenizer, and then centrifuged at 700 g for 10 min in 4C to collect the cytosol fraction in the supernatant. The cytosol fraction was further centrifuged at 10,000 g for 15 min to collect the mitochondria fraction in the pellet. Same amounts of SDS-reduced cytosol or mitochondria fractions were run in western blot, and detected with antibody 15-7, HSPA9 and GAPDH (Cell Signaling Technology, #97166). A separate sample gel was silver stained and cut around 60kd for mass spectrometry analysis.

### Histopathology image analysis

The scanned resected PDAC histopathologic images were opened with ImageScope (Leica), and lymphocytes aggregates with bigger than 0.01 mm^2 size were counted and exported for size measurement using image J.

### Statistical analysis

Statistical analysis was performed using Graphpad Prism 5. The plasma IgG data was first tested for normality distribution in Graphpad. Mann-Whitney t-test was used for a two-group comparison of non-normal distribution data, Kruskal-Wallis ANOVA test was used for comparisons of three or more groups of non-normal distribution data. Fisher’s test was used for demographic factors comparison, and Pearson’s correlation was used for correlation studies. P<0.05 was used for statistical significance.

## List of Supplementary Materials

Fig. S1 Characterization of primary PDAC immune cells microenvironment by scRNA-Seq

Fig. S2. Presence of TLS like structures in PDAC.

Fig. S3 Additional Ig sequences evolution trees from PCs in PDAC

Fig. S4 Antibodies staining atlas on MiaPaca2 cell line.

Fig. S5 Examples of antibodies staining in PDAC and non-cancer cell lines

Fig. S6 Characterization of antibodies 8-3, 19-3 and 15-7

Fig. S7 Comparison of plasma IgG titers to F-actin, RUVBL2 and HSPD1 between healthy donors and PDAC patients.

Table S1 Human PDAC samples information for scRNA-Seq studies.

Table S2 List of antibodies synthesized and screening results.

Table S3 Human samples information for plasma studies.

## Acknowledgments

We thank Sharon Fox and Cristina Valente and other members from Northwell Health Biospecimen Repository in helping coordinate the human sample collections. We thank Ledong Wan from Dr. Krainer lab at CSHL in providing the HPDE cell line. We thank Dr. Semir Bayez of CSHL for his thoughtful input in this project. We thank members of Fearon lab for their thoughtful discussions of this project and help review of this manuscript. This project was funded by Lustgarten Foundation and Simons Foundation awarded to Dr. Douglas Fearon.

## Author contributions

Conceptualization: MY, DF

Methodology: MY, JP, JY, DP, PC, YZ, BH, HP, AH, DK, KP, AR, DS, MW, DT

Investigation: MY, SS, PM,

Visualization: MY, JP, YZ, BH

Funding acquisition: DF

Project administration: MY

Supervision: DF

Writing – original draft: MY, DF

Writing – review & editing: YZ, PM, KP, AR

## Data and materials availability

The sequencing data from this study will be deposited into the NCBI database. Public codes/software was used for the data analysis as described in the paper. The codes for sequencing data analysis and experimental material can be accessed through communication with the corresponding authors.

## Supplementary materials

### Supplementary figures

**Fig. S1.**
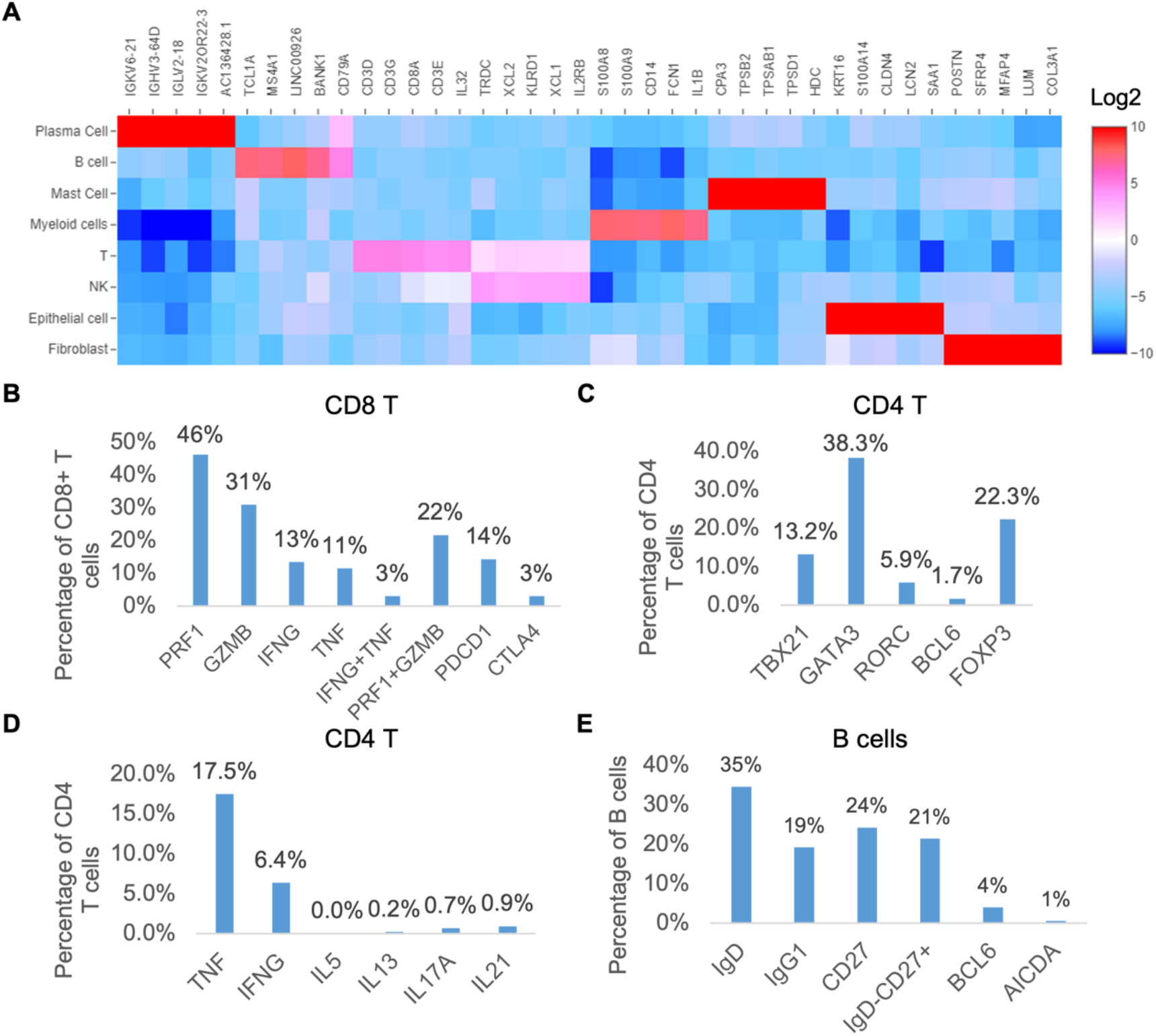
Characterization of primary PDAC immune cells microenvironment by scRNA-Seq. **(A)** Heatmap of the top five most upgraded genes in each cell cluster defined in Figure 1A. Color scale is log2 fold changes. (**B-E**) The percentage of cells expressing selected gene markers in the clusters of CD8 T (**B**), CD4 (**C, D**) and B cells (**E**).

**Fig. S2.**
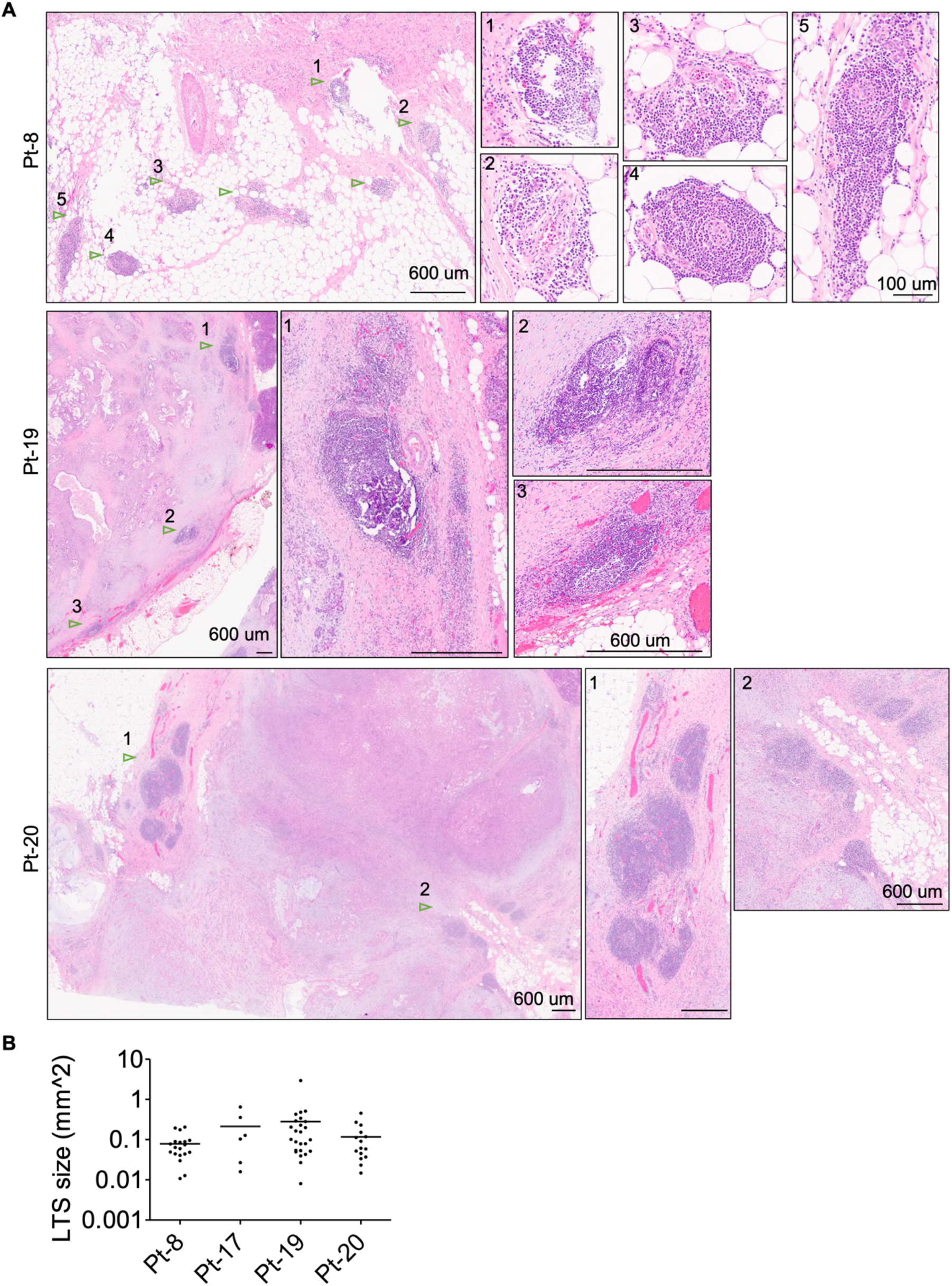
Presence of TLS like structures in PDAC. **(A)** Examples of TLS like structures in patient 8, 19 and 20 (TLS indicated by arrowhead and labelled by numbers). The zoom in images were shown in the right. Scale bar is shown in figures. (**B**) Quantification and size distribution of the TLS in the four resected PDAC samples. Only TLS with size bigger than 0.01 mm^2 were analyzed.

**Fig. S3.**
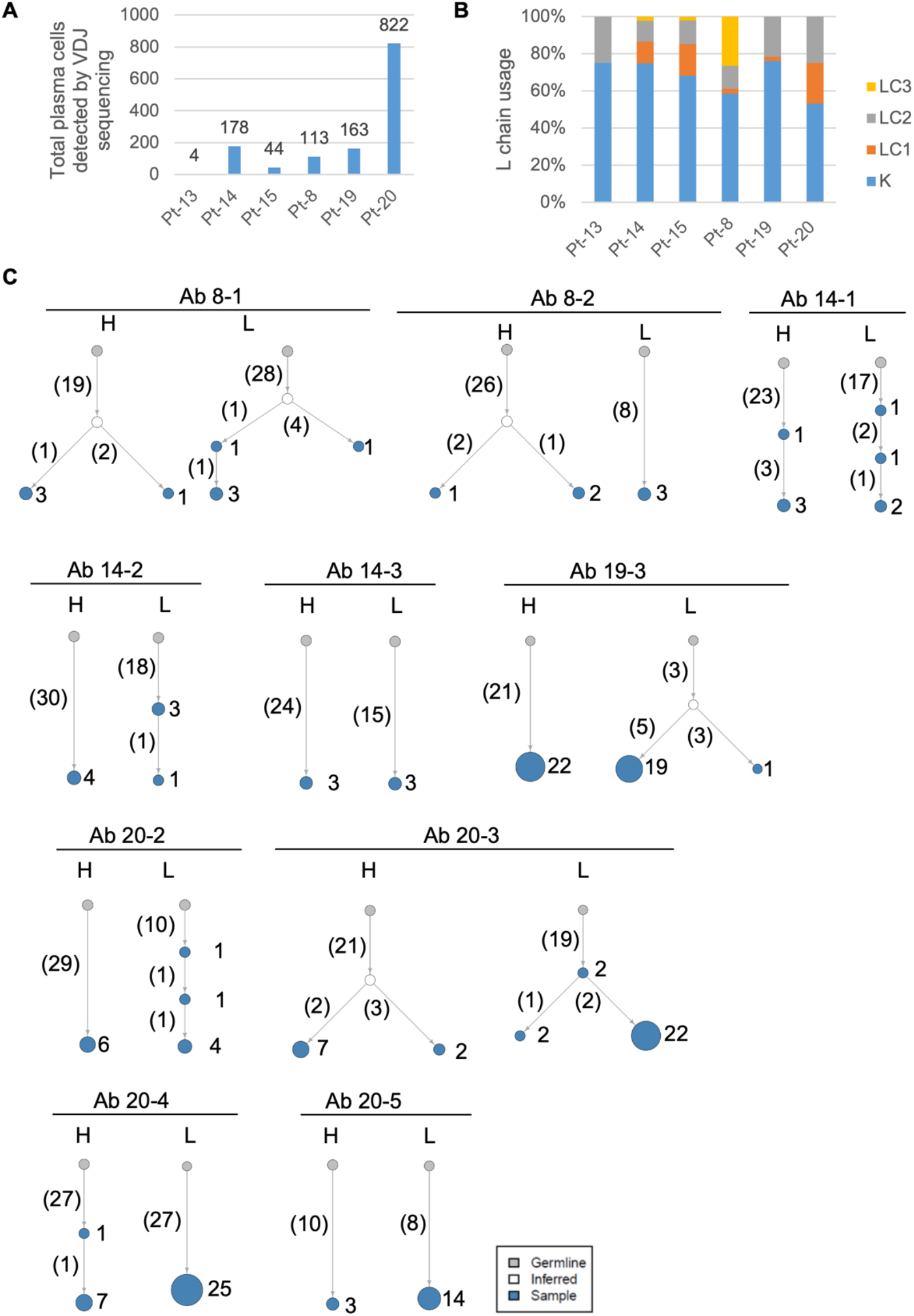
Additional Ig sequences evolution trees from PCs in PDAC. Additional examples of antibody lineage evolution among top expanded PC clones screened in this study. Clone size is indicated by the size of the node (not scaled) and labeled by numbers on the right. The clone size includes both paired and unpaired chains. Number of somatic mutations in combined V(D)J regions is shown in parenthesis.

**Fig. S4.**
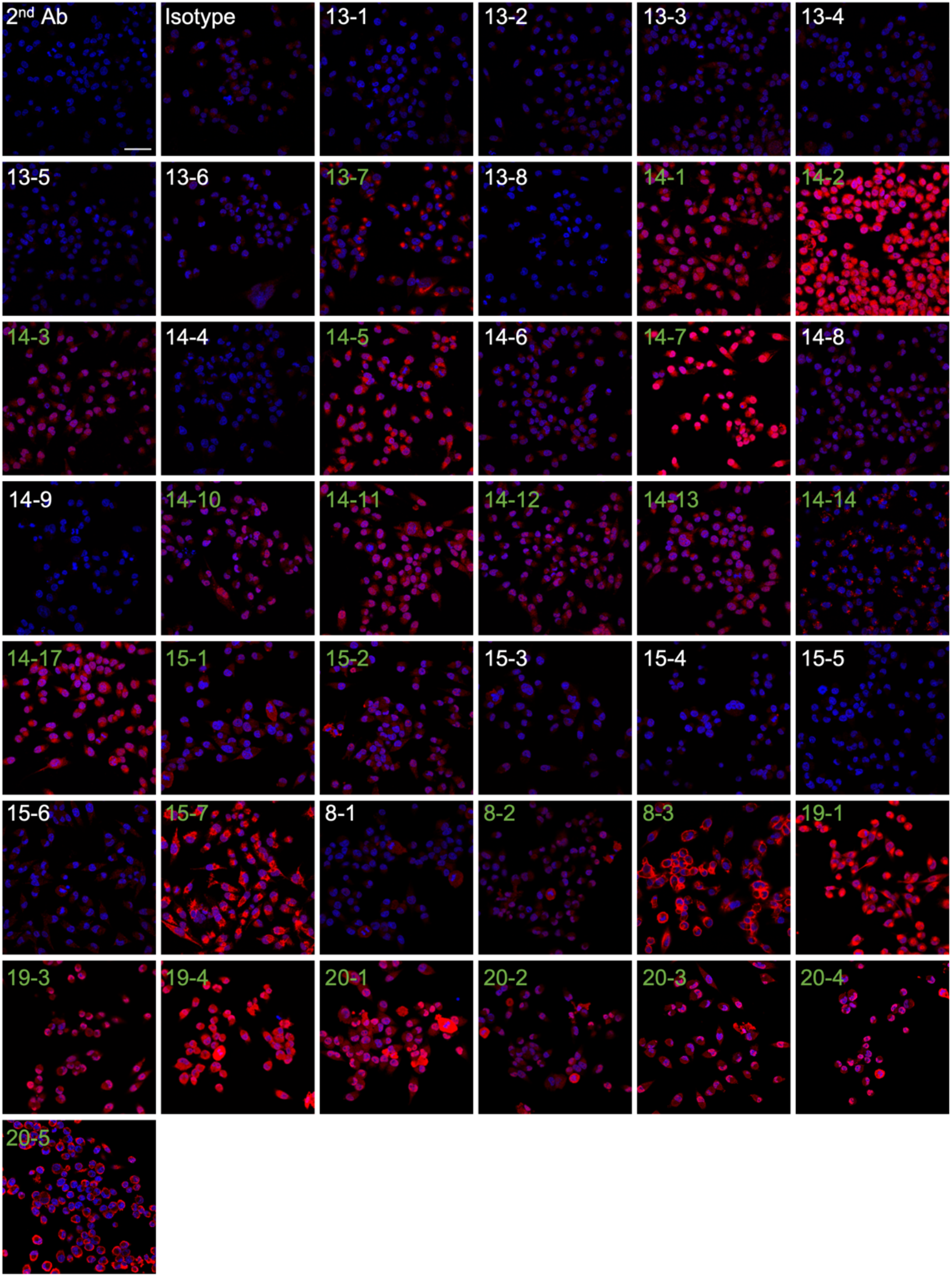
Antibodies staining atlas on MiaPaca2 cell line. MiaPaca2 cells were stained with each antibody (red) and co-stained with DAPI (blue). Secondary antibody only (2^nd^ Ab) and isotype IgG were used as control. Antibodies with positive staining are labeled green, and antibodies with negative staining are labeled white. All images were acquired using the same confocal setting for isotype control staining, and the gain on the query antibody was reduced if the images were saturated. Scale bar is 50um.

**Fig. S5.**
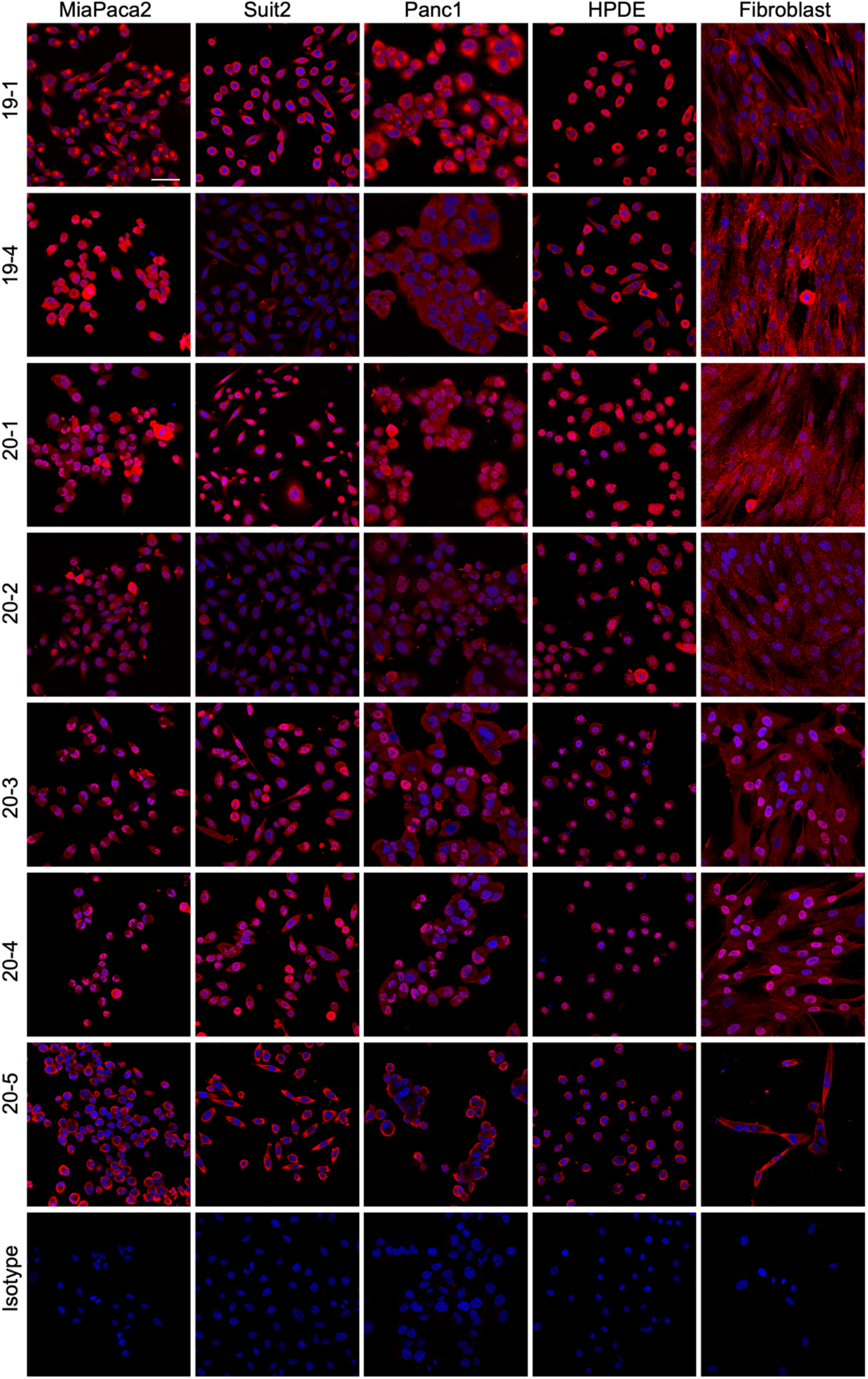
Examples of antibodies staining in PDAC and non-cancer cell lines. Cells were stained with each antibody (red) and co-stained with DAPI (blue). Scale bar is 50um.

**Fig. S6.**
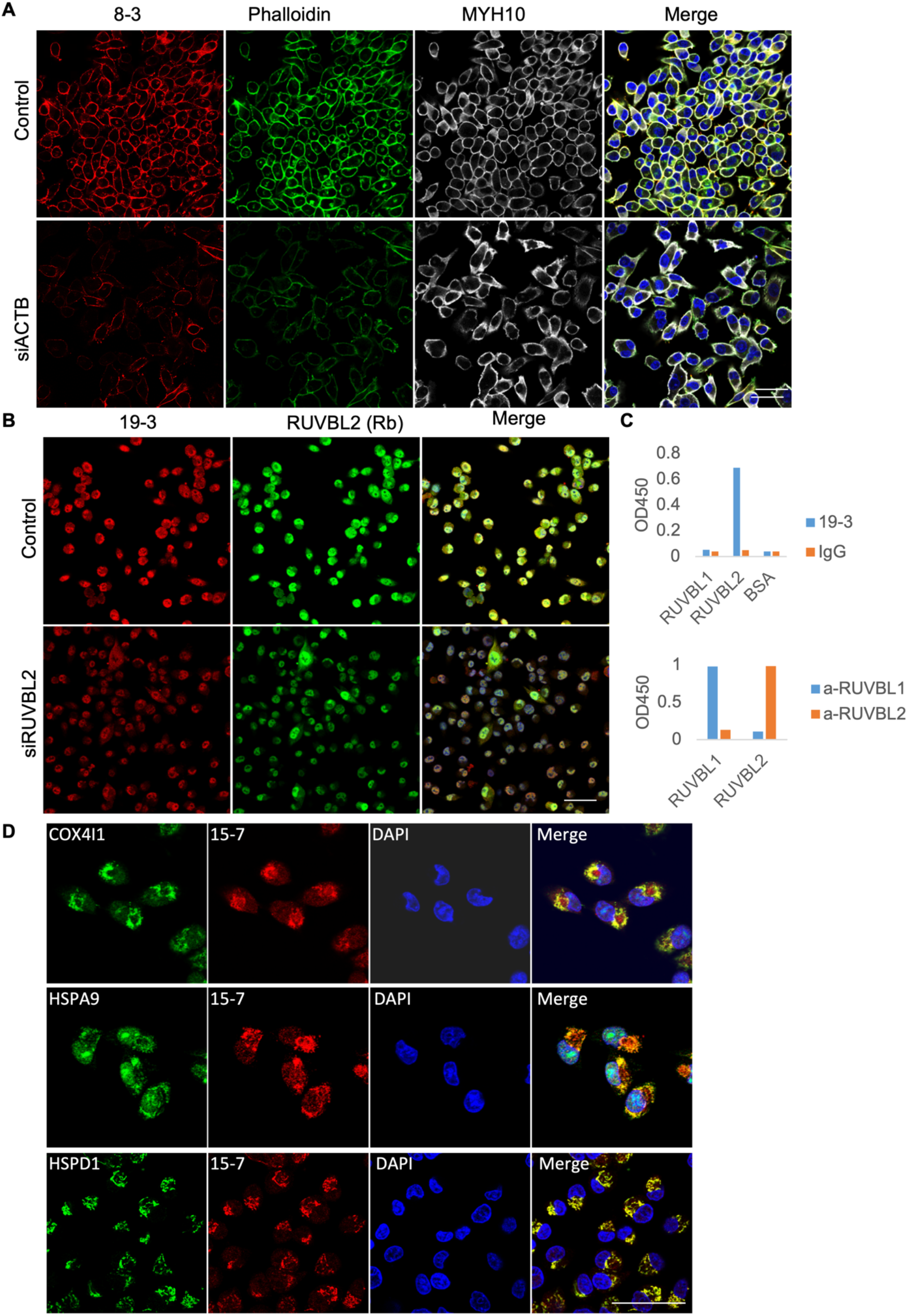
Characterization of antibodies 8-3, 19-3 and 15-7. (**A**) 8-3 staining after siRNA knockdown of *ACTB* in MiaPaca2 cells, co-stained with phalloidin, MYH10 antibody and DAPI. (**B**) Measurement of 19-3 binding to recombinant RUVBL1 and RUBL2 proteins by ELISA. Positive controls using polyclonal rabbit antibodies anti-RUVBL1 (a-RUVBL1) or anti-RUVBL2 (a-RUVBL2). (**C**) 19-3 staining after siRNA knockdown of *RUVBL2* in MiaPaca2 cells, co-stained with anti-RUVBL2 rabbit antibody and DAPI. (**D**) Co-staining of 15-7 with mitochondria markers COX4I1, HSPA9 and HSPD1 in MiaPaca2 cells. Scale bars in A, C and D are 50um.

**Fig. S7.**
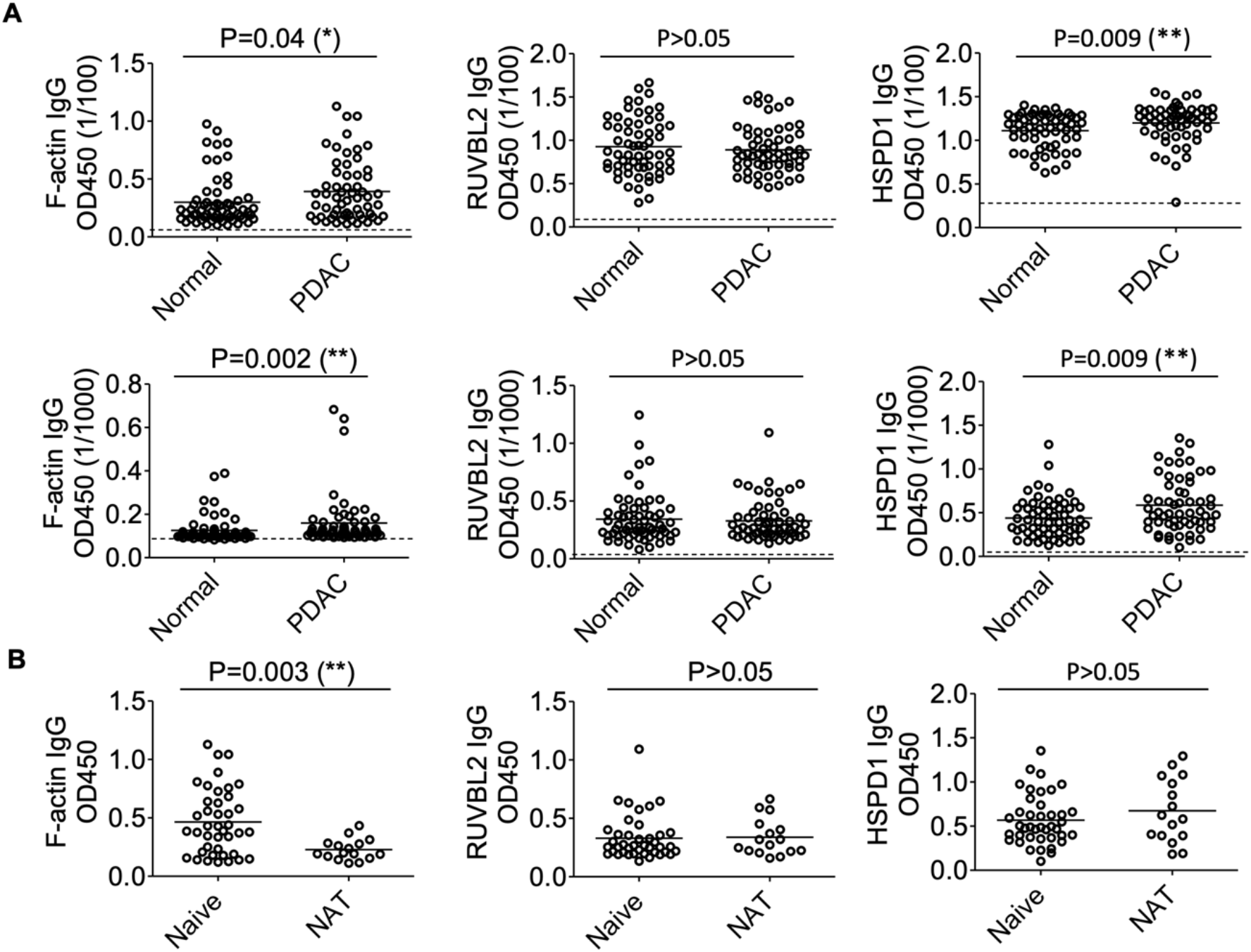
Comparison of plasma IgG titers to F-actin, RUVBL2 and HSPD1 between healthy donors and PDAC patients. (**A**) Comparison of IgG titer to F-actin, RUVBL2 and HSPD1 of plasma samples diluted at 1/100 or 1/1000 by ELISA. Background signals from secondary antibody only are shown in a dashed line. (**B**) Comparison of IgG response to F-actin, RUVBL2 and HSPD1 between naïve treated (naive) and neoadjuvant treated (NAT) PDAC patients. The non-parametric t-test is used for comparisons.

**Table S1.** Human PDAC samples information for scRNA-Seq studies.

|  |  |  |  |  |  |  |  |
| --- | --- | --- | --- | --- | --- | --- | --- |
| **Patient ID** | Pt-8 | Pt-17 | Pt-19 | Pt-20 | Pt-13 | Pt-14 | Pt-15 |
| **Diagnosis** | PDAC | PDAC | PDAC | PDAC | PDAC | PDAC | PDAC |
| **Collection method** | resection | resection | resection | resection | FNA | FNA | FNA |
| **Age** | 58 | 75 | 71 | 67 | 40 | 89 | 53 |
| **Gender** | Male | Male | Female | Female | Female | Male | Female |
| **Race** | Caucasian | Caucasian | African-American | Asian | Latino/Hispanic | Caucasian | African American |
| **Neoadjuvant therapy** | No | No | No | No | No | No | No |
| **TNM stage** | pT2N2Mx | pT2N1Mx | pT2N1 | pT2N0 | NA | NA | NA |
| **Tumor size (cm)** | 2.2*1.5*1 | 2.7*2.6*2.5 | 2.7*2.3*2.2 | 2.2*1.7*0.8 | NA | NA | NA |
| **MSI status** | MSS | MSS | MSS | MSS | NA | NA | NA |
| **Tumor organoid available** | Yes | No | No | Yes | Yes | Yes | No |
| **Tumor section available** | Yes | Yes | Yes | Yes | No | No | No |
| **Gene Expressing Sequencing** | Yes | Yes | Yes | Yes | Yes | Yes | Failed |
| **Ig sequencing** | Yes | Yes (low B and PC cells) | Yes | Yes | Yes | Yes | Yes |

Abbreviations: FNA, fine needle aspiration; MSI, microsatellite instable; MSS, microsatellite stable; NA, not available/appliable

**Table S2.** List of antibodies synthesized and screening results.

| No | Patient | Ab-name | Clone size | Cell type | Isotype | Light chain | Binding reactivity | Antigen location | Antigen identity | Synthesize format |
| --- | --- | --- | --- | --- | --- | --- | --- | --- | --- | --- |
| 1 | Pt-8 | 8-1 | 4 | PC | IgG1 | IgK | Negative | NA |  | IgG4-His |
| 2 | Pt-8 | 8-2 | 3 | PC | IgA1 | IgK | Positive | nuclei enriched | Unknown | IgG4-His |
| 3 | Pt-8 | 8-3 | 20 | PC | IgG2 | IgLC3 | Positive | cyto | F-actin | IgG4-His |
| 4 | Pt-19 | 19-1 | 46 | PC | IgG1 | IgK | Positive | cyto enriched | Unknown | IgG4-His |
| 5 | Pt-19 | 19-3 | 23 | PC | IgG1 | IgK | Positive | nuclei enriched | RUVBL2 | IgG4-His |
| 6 | Pt-19 | 19-4 | 3 | PC | IgA2 | IgK | Positive | cyto/nuclei even | Unknown | IgG4-His |
| 7 | Pt-20 | 20-1 | 25 | PC | IgG2 | IgLC2 | positive | nuclei enriched | Unknown | IgG4-His |
| 8 | Pt-20 | 20-2 | 6 | PC | IgG2 | IgLC1 | positive | nuclei enriched | Unknown | IgG4-His |
| 9 | Pt-20 | 20-3 | 26 | PC | IgG2 | IgLC1 | positive | nuclei enriched | Unknown | IgG4-His |
| 10 | Pt-20 | 20-4 | 25 | PC | IgG2 | IgK | positive | nuclei enriched | Unknown | IgG4-His |
| 11 | Pt-20 | 20-5 | 14 | PC | IgG2 | IgK | positive | cyto | Unknown | IgG4-His |
| 12 | Pt-13 | 13-1 | 1 | PC | IgG1 | IgK | Negative | NA |  | IgG1 |
| 13 | Pt-13 | 13-2 | 1 | PC | IgG1 | IgLC2 | Negative | NA |  | IgG1 |
| 14 | Pt-13 | 13-3 | 1 | PC | IgA1 | IgK | Negative | NA |  | IgG1 |
| 15 | Pt-13 | 13-4 | 1 | PC | IgG1 | IgK | Negative | NA |  | IgG1 |
| 16 | Pt-13 | 13-5 | 1 | PC | IgA1 | IgK | Negative | NA |  | IgG1 |
| 17 | Pt-13 | 13-6 | 1 | GC-B | IgG1 | IgK | Negative | NA |  | IgG1 |
| 18 | Pt-13 | 13-7 | 1 | GC-B | IgG3 | IgLC2 | Positive | cyto | Unknown | IgG1 |
| 19 | Pt-13 | 13-8 | 2 | B | IgA1 | IgLC2 | Negative | NA |  | IgG1 |
| 20 | Pt-14 | 14-1 | 4 | PC | IgA2 | IgK | Positive | cyto/nuclei even | Unknown | IgG1 |
| 21 | Pt-14 | 14-2 | 4 | PC | IgA2 | IgK | Positive | cyto/nuclei even | Unknown | IgG1 |
| 22 | Pt-14 | 14-3 | 3 | PC | IgA2 | IgK | Positive | nuclei enriched | Unknown | IgG1 |
| 23 | Pt-14 | 14-4 | 2 | PC | IgM | IgK | Negative | NA |  | IgG1 |
| 24 | Pt-14 | 14-5 | 2 | PC | IgA1 | IgK | Positive | cyto enriched | Unknown | IgG1 |
| 25 | Pt-14 | 14-6 | 2 | PC | IgA2 | IgK | Negative | NA |  | IgG1 |
| 26 | Pt-14 | 14-7 | 2 | PC | IgA2 | IgK | Positive | nuclei enriched | Unknown | IgG1 |
| 27 | Pt-14 | 14-8 | 2 | PC | IgA2 | IgLC3 | Negative | NA |  | IgG1 |
| 28 | Pt-14 | 14-9 | 1 | PC | IgM | IgK | Negative | NA |  | IgG1 |
| 29 | Pt-14 | 14-10 | 1 | PC | IgM | IgLC2 | Positive | nuclei enriched | Unknown | IgG1 |
| 30 | Pt-14 | 14-11 | 1 | PC | IgA2 | IgLC1 | Positive | nuclei enriched | Unknown | IgG1 |
| 31 | Pt-14 | 14-12 | 1 | PC | IgG1 | IgK | Positive | nuclei enriched | Unknown | IgG1 |
| 32 | Pt-14 | 14-13 | 1 | PC | IgA2 | IgK | Positive | cyto/nuclei<br>even | Unknown | IgG1 |
| 33 | Pt-14 | 14-14 | 1 | PC | IgA2 | IgK | Positive | cyto | Unknown | IgG1 |
| 34 | Pt-14 | 14-17 | 1 | PC | IgG3 | IgK | Positive | cyto<br>enriched | Unknown | IgG1 |
| 35 | Pt-15 | 15-1 | 3 | B | IgA2 | IgK | Positive | cyto/nuclei<br>even | Unknown | IgG1 |
| 36 | Pt-15 | 15-2 | 3 | B | IgA1 | IgK | Positive | cyto/nuclei<br>even | Unknown | IgG1 |
| 37 | Pt-15 | 15-3 | 2 | PC | IgM | IgKC | Negative | NA |  | IgG1 |
| 38 | Pt-15 | 15-4 | 2 | PC | IgA1 | IgK | Negative | NA |  | IgG1 |
| 39 | Pt-15 | 15-5 | 2 | PC | IgA2 | IgK | Negative | NA |  | IgG1 |
| 40 | Pt-15 | 15-6 | 2 | B | IgA2 | IgLC1 | Negative | NA |  | IgG1 |
| 41 | Pt-15 | 15-7 | 1 | PC | IgG1 | IgK | Positive | cyto | HSPD1 | IgG1 |

Abbreviations: PC-plasma cells, B- B cells, GC-B: Germinal center B cells, cyto-cytoplasmic
Light orange color indicates the three antibodies that have antigens identified.

**Table S3.**
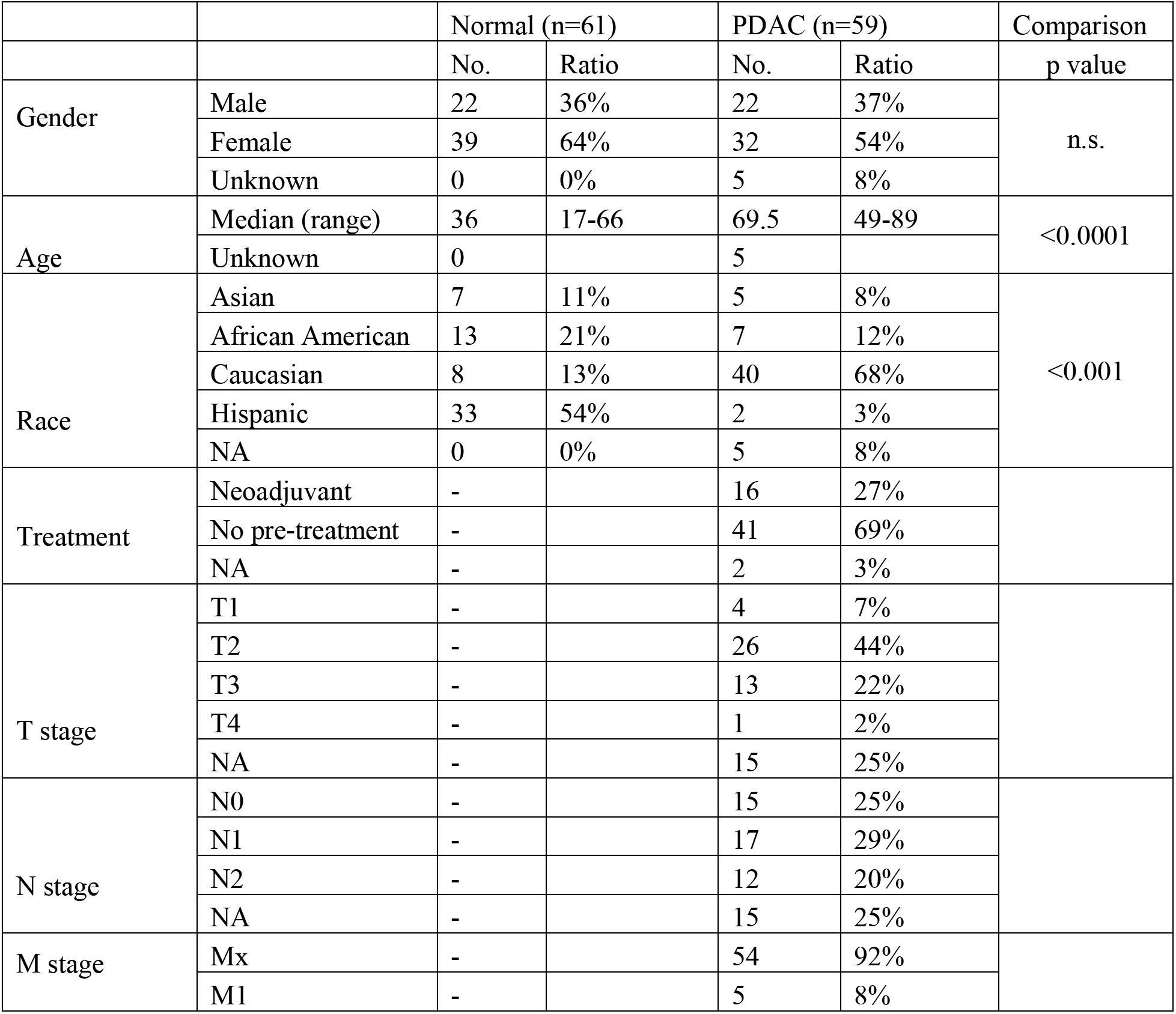
Human samples information for plasma studies.

## References and Notes

1. A. S. Bear, R. H. Vonderheide, M. H. O’Hara, Challenges and Opportunities for Pancreatic Cancer Immunotherapy. Cancer Cell 38, 788-802 (2020).

2. N. Hiraoka, Y. Ino, R. Yamazaki-Itoh, Y. Kanai, T. Kosuge, K. Shimada, Intratumoral tertiary lymphoid organ is a favourable prognosticator in patients with pancreatic cancer. Br J Cancer 112, 1782-1790 (2015).

3. G. F. Castino, N. Cortese, G. Capretti, S. Serio, G. Di Caro, R. Mineri, E. Magrini, F. Grizzi, P. Cappello, F. Novelli, P. Spaggiari, M. Roncalli, C. Ridolfi, F. Gavazzi, A. Zerbi, P. Allavena, F. Marchesi, Spatial distribution of B cells predicts prognosis in human pancreatic adenocarcinoma. Oncoimmunology 5, e1085147 (2016).

4. J. G. A, V. Rajamanickam, C. Bui, B. Bernard, J. Pucilowska, C. Ballesteros-Merino, M. Schmidt, K. McCarty, M. Philips, B. Piening, C. Dubay, T. Medler, P. Newell, P. Hansen, E. Tran, E. Tang, C. Bifulco, M. Crittenden, M. Gough, K. H. Young, Germinal center reactions in tertiary lymphoid structures associate with neoantigen burden, humoral immunity and long-term survivorship in pancreatic cancer. Oncoimmunology 10, 1900635 (2021).

5. D. Biasci, M. Smoragiewicz, C. M. Connell, Z. Wang, Y. Gao, J. E. D. Thaventhiran, B. Basu, L. Magiera, T. I. Johnson, L. Bax, A. Gopinathan, C. Isherwood, F. A. Gallagher, M. Pawula, I. Hudecova, D. Gale, N. Rosenfeld, P. Barmpounakis, E. C. Popa, R. Brais, E. Godfrey, F. Mir, F. M. Richards, D. T. Fearon, T. Janowitz, D. I. Jodrell, CXCR4 inhibition in human pancreatic and colorectal cancers induces an integrated immune response. Proc Natl Acad Sci U S A 117, 28960-28970 (2020).

6. V. P. Balachandran, M. Luksza, J. N. Zhao, V. Makarov, J. A. Moral, R. Remark, B. Herbst, G. Askan, U. Bhanot, Y. Senbabaoglu, D. K. Wells, C. I. O. Cary, O. Grbovic-Huezo, M. Attiyeh, B. Medina, J. Zhang, J. Loo, J. Saglimbeni, M. Abu-Akeel, R. Zappasodi, N. Riaz, M. Smoragiewicz, Z. L. Kelley, O. Basturk, I. Australian Pancreatic Cancer Genome, R. Garvan Institute of Medical, H. Prince of Wales, H. Royal North Shore, G. University of, H. St Vincent’s, Q. B. M. R. Institute, C. f. C. R. University of Melbourne, I. f. M. B. University of Queensland, H. Bankstown, H. Liverpool, C. O. B. L. Royal Prince Alfred Hospital, H. Westmead, H. Fremantle, H. St John of God, H. Royal Adelaide, C. Flinders Medical, P. Envoi, H. Princess Alexandria, H. Austin, I. Johns Hopkins Medical, A. R.-N. C. f. A. R. o. Cancer, M. Gonen, A. J. Levine, P. J. Allen, D. T. Fearon, M. Merad, S. Gnjatic, C. A. Iacobuzio-Donahue, J. D. Wolchok, R. P. DeMatteo, T. A. Chan, B. D. Greenbaum, T. Merghoub, S. D. Leach, Identification of unique neoantigen qualities in long-term survivors of pancreatic cancer. Nature 551, 512-516 (2017).

7. N. G. Kooreman, Y. Kim, P. E. de Almeida, V. Termglinchan, S. Diecke, N. Y. Shao, T. T. Wei, H. Yi, D. Dey, R. Nelakanti, T. P. Brouwer, D. T. Paik, I. Sagiv-Barfi, A. Han, P. H. A. Quax, J. F. Hamming, R. Levy, M. M. Davis, J. C. Wu, Autologous iPSC-Based Vaccines Elicit Anti-tumor Responses In Vivo. Cell Stem Cell 22, 501-513 e507 (2018).

8. X. Ouyang, Y. Liu, Y. Zhou, J. Guo, T. T. Wei, C. Liu, B. Lee, B. Chen, A. Zhang, K. M. Casey, L. Wang, N. G. Kooreman, A. Habtezion, E. G. Engleman, J. C. Wu, Antitumor effects of iPSC-based cancer vaccine in pancreatic cancer. Stem Cell Reports 16, 1468-1477 (2021).

9. R. Di Niro, L. Mesin, N. Y. Zheng, J. Stamnaes, M. Morrissey, J. H. Lee, M. Huang, R. Iversen, M. F. du Pre, S. W. Qiao, K. E. Lundin, P. C. Wilson, L. M. Sollid, High abundance of plasma cells secreting transglutaminase 2-specific IgA autoantibodies with limited somatic hypermutation in celiac disease intestinal lesions. Nat Med 18, 441-445 (2012).

10. C. Luchini, L. A. A. Brosens, L. D. Wood, D. Chatterjee, J. I. Shin, C. Sciammarella, G. Fiadone, G. Malleo, R. Salvia, V. Kryklyva, M. L. Piredda, L. Cheng, R. T. Lawlor, V. Adsay, A. Scarpa, Comprehensive characterisation of pancreatic ductal adenocarcinoma with microsatellite instability: histology, molecular pathology and clinical implications. Gut 70, 148-156 (2021).

11. E. Elyada, M. Bolisetty, P. Laise, W. F. Flynn, E. T. Courtois, R. A. Burkhart, J. A. Teinor, P. Belleau, G. Biffi, M. S. Lucito, S. Sivajothi, T. D. Armstrong, D. D. Engle, K. H. Yu, Y. Hao, C. L. Wolfgang, Y. Park, J. Preall, E. M. Jaffee, A. Califano, P. Robson, D. A. Tuveson, Cross-Species Single-Cell Analysis of Pancreatic Ductal Adenocarcinoma Reveals Antigen-Presenting Cancer-Associated Fibroblasts. Cancer Discov 9, 1102-1123 (2019).

12. J. DeFalco, M. Harbell, A. Manning-Bog, G. Baia, A. Scholz, B. Millare, M. Sumi, D. Zhang, F. Chu, C. Dowd, P. Zuno-Mitchell, D. Kim, Y. Leung, S. Jiang, X. Tang, K. S. Williamson, X. Chen, S. M. Carroll, G. Espiritu Santo, N. Haaser, N. Nguyen, E. Giladi, D. Minor, Y. C. Tan, J. B. Sokolove, L. Steinman, T. A. Serafini, G. Cavet, N. M. Greenberg, J. Glanville, W. Volkmuth, D. E. Emerling, W. H. Robinson, Non-progressing cancer patients have persistent B cell responses expressing shared antibody paratopes that target public tumor antigens. Clin Immunol 187, 37-45 (2018).

13. R. D. Mazor, N. Nathan, A. Gilboa, L. Stoler-Barak, L. Moss, I. Solomonov, A. Hanuna, Y. Divinsky, M. D. Shmueli, H. Hezroni, I. Zaretsky, M. Mor, O. Golani, G. Sabah, A. Jakobson-Setton, N. Yanichkin, M. Feinmesser, D. Tsoref, L. Salman, E. Yeoshoua, E. Peretz, I. Erlich, N. M. Cohen, J. M. Gershoni, N. Freund, Y. Merbl, G. Yaari, R. Eitan, I. Sagi, Z. Shulman, Tumor-reactive antibodies evolve from non-binding and autoreactive precursors. Cell 185, 1208-1222 e1221 (2022).

14. H. Wardemann, S. Yurasov, A. Schaefer, J. W. Young, E. Meffre, M. C. Nussenzweig, Predominant autoantibody production by early human B cell precursors. Science 301, 1374-1377 (2003).

15. A. Granito, L. Muratori, P. Muratori, G. Pappas, M. Guidi, F. Cassani, U. Volta, A. Ferri, M. Lenzi, F. B. Bianchi, Antibodies to filamentous actin (F-actin) in type 1 autoimmune hepatitis. J Clin Pathol 59, 280-284 (2006).

16. K. Kaji, N. Fertig, T. A. Medsger, Jr., T. Satoh, K. Hoshino, Y. Hamaguchi, M. Hasegawa, M. Lucas, A. Schnure, F. Ogawa, S. Sato, K. Takehara, M. Fujimoto, M. Kuwana, Autoantibodies to RuvBL1 and RuvBL2: a novel systemic sclerosis-related antibody associated with diffuse cutaneous and skeletal muscle involvement. Arthritis Care Res (Hoboken) 66, 575-584 (2014).

17. G. Schett, Q. Xu, A. Amberger, R. Van der Zee, H. Recheis, J. Willeit, G. Wick, Autoantibodies against heat shock protein 60 mediate endothelial cytotoxicity. J Clin Invest 96, 2569-2577 (1995).

18. M. H. Hansen, H. Nielsen, H. J. Ditzel, The tumor-infiltrating B cell response in medullary breast cancer is oligoclonal and directed against the autoantigen actin exposed on the surface of apoptotic cancer cells. Proc Natl Acad Sci U S A 98, 12659-12664 (2001).

19. Y. He, Y. Wu, Z. Mou, W. Li, L. Zou, T. Fu, A. Zhang, D. Xiang, H. Xiao, X. Wang, Proteomics-based identification of HSP60 as a tumor-associated antigen in colorectal cancer. Proteomics Clin Appl 1, 336-342 (2007).

20. E. Giampazolias, O. Schulz, K. H. J. Lim, N. C. Rogers, P. Chakravarty, N. Srinivasan, O. Gordon, A. Cardoso, M. D. Buck, E. Z. Poirier, J. Canton, S. Zelenay, S. Sammicheli, N. Moncaut, S. Varsani-Brown, I. Rosewell, C. Reis e Sousa, Secreted gelsolin inhibits DNGR-1-dependent cross-presentation and cancer immunity. Cell 184, 4016-4031 e4022 (2021).

21. S. F. Boj, C. I. Hwang, L. A. Baker, Chio, II, D. D. Engle, V. Corbo, M. Jager, M. Ponz-Sarvise, H. Tiriac, M. S. Spector, A. Gracanin, T. Oni, K. H. Yu, R. van Boxtel, M. Huch, K. D. Rivera, J. P. Wilson, M. E. Feigin, D. Ohlund, A. Handly-Santana, C. M. Ardito-Abraham, M. Ludwig, E. Elyada, B. Alagesan, G. Biffi, G. N. Yordanov, B. Delcuze, B. Creighton, K. Wright, Y. Park, F. H. Morsink, I. Q. Molenaar, I. H. Borel Rinkes, E. Cuppen, Y. Hao, Y. Jin, I. J. Nijman, C. Iacobuzio-Donahue, S. D. Leach, D. J. Pappin, M. Hammell, D. S. Klimstra, O. Basturk, R. H. Hruban, G. J. Offerhaus, R. G. Vries, H. Clevers, D. A. Tuveson, Organoid models of human and mouse ductal pancreatic cancer. Cell 160, 324-338 (2015).

22. N. T. Gupta, J. A. Vander Heiden, M. Uduman, D. Gadala-Maria, G. Yaari, S. H. Kleinstein, Change-O: a toolkit for analyzing large-scale B cell immunoglobulin repertoire sequencing data. Bioinformatics 31, 3356-3358 (2015).

